# Direct determination and differentiation of carbapenemases of *A. baumannii* from uncultured tracheal samples

**DOI:** 10.1101/347047

**Authors:** CH. Swathi, S. Sukanya, V. Lakshmi, K.S. Ratnakar, V. Sritharan

**Author notes:** Address correspondence to V. Sritharan. Present address: V. Lakshmi, Kamineni Academy of Medical Sciences and Research Center, LN nagar, Hyderabad.

## Abstract

The emergence of multiple carbapenemases and the consequent multi drug resistance in bacteria constitute a grave concern in the management of critical care patients and also in community acquired infections. Detection of carbapenamase activity helps to understand the possible mechanism(s) of carbapenem resistance in the microorganism. Identification of carbapenemases is currently being done by various phenotypic methods and molecular methods. However, innovative biochemical and spectrophotometric methods are desirable as they will be easy to perform, affordable and rapid. Recently a novel chromogenic method called CarbaNP test was introduced to screen for carbapenemases in clinical isolates of gram negative pathogens. We adopted this assay: (i) to detect the total carbapenemase activity (ii) to measure the relative rates of hydrolysis of imipenem by class A, B and D carbapenemases with inhibitors (iii) to confirm the genotype by PCR and (iv)for direct differential detection of various carbapenemases in uncultured clinical sample like tracheal aspirate. The study included 75 culture isolates and 153 purulent tracheal aspirates. All isolates were screened by our optimized protocol and also genotyped by PCR. This adopted assay showed good sensitivity and correlation with conventional phenotyping and genotyping. Our protocol offers the fastest way to identify the pathogen by PCR but also its carbapenemase profile directly from uncultured clinical samples in less than 4h. Our protocol is currently being validated on other types of clinical specimens in our laboratory.

## Introduction

The emergence of multidrug resistance (MDR) among gram negative pathogens is severely hampering the management of infections in the hospitals as well as in the community. Carbapenemases hydrolyse beta lactam antibiotics and all types of carbapenems. Their production is one of the most important traits which confers resistance to beta lactam antibiotics including carbapenems [1, 2]. In recent years laboratory detection of carbapenemase producers has become more challenging due to the emergence of diverse carbapenemases [3, 4]. Rapid detection and carbapenem sensitivity profiling of these pathogens have become imperative for successful management of infections and also to control dissemination. Antimicrobial susceptibility tests are usually performed on agar plates or in automated systems. Carbapenemases are not the sole cause of resistance, other mechanisms such as porin loss or increased efflux pump activity in the bacterial membrane may also influence the drug sensitivity [5, 6, 7 & 8]. Therefore a definite method is required to distinguish enzymatic and non-enzymatic resistance mechanisms for effective management of infection control and epidemic outbreaks [2, 9]. Phenotypic methods like Modified Hodge Test (MHT) and disk diffusion tests with inhibitors are commonly used in microbiology laboratory to test the carbapenemase activity of isolated bacterial cultures. However, these methods show lower specificity and sensitivity compared to molecular methods [10]. Identification of the resistance gene is considered as gold standard for the confirmation of drug resistance, though they are economically and technically beyond the reach for many clinical laboratories [3, 5]. Thus, there has been an unmet need for a simple, affordable and yet direct demonstration of carbapenamase activity in any given bacterial pathogen. To fill this important gap, Nordmann group designed and developed a novel yet simple phenotypic method called CarbaNP test, for early detection of carbapenemases in *Enterobacteriaceae* [2]. This phenotypic assay is based on the change in pH caused due to hydrolysis of imipenem by carbapenamase(s) present in the cell free bacterial lysate. Phenol red is used as the indicator to monitor this change in pH, which turns from red (alkaline) to yellow (acid) [2, 3, 11 and 12]. This test has become a boon for rapid identification of carbapenemase production in gram negative bacterial pathogens, considering the difficulties in the interpretation of results and false positivity with various other phenotypic methods. CarbaNP is a huge improvement over other phenotypic methods with almost 100% sensitivity and its specificity is comparable to molecular methods [2]. Several modifications have been made to this assay in order to improve its performance characteristics and avoid false negatives [10, 13 and 14]. Different classes of carbapenemases have been reported in gram negative pathogens. There are three major classes of carbapenemases, grouped according to their amino acid composition and identity [15]. Based on the action of beta-lactamase inhibitors (tazobactum, clavulanic acid and divalent chelator such as EDTA) CarbaNP assay has been modified to identify different classes of carbapenemases [12] in gram negative bacterial isolates. Later, “Blue-Carba”, was designed using bromothymol blue as an indicator for direct colony approach instead of bacterial extracts [16]. Originally CarbaNP assay was employed for members of *Enterobacteriaceae* isolates, later it was applied to isolates of *Acinetobacter* sp. with minor changes in inoculum size and lysis conditions [17]. The main objective of this study is to adopt and evaluate this CarbaNP assay for the rapid identification and differentiation of various classes of carbapenemases on clinical isolates of *A.baumannii* and extend it to uncultured clinical specimens. In this study we adopted CarbaNP test of Nordmann’s with some modifications in sample processing and assay protocols. In addition, we validated our protocol directly on uncultured tracheal aspirates also. We included specific inhibitors for differential measurement of carbapenemases belonging to Class A, B, and D. We believe that these modifications simplify the assay for application to both culture isolates and directly to clinical samples also. These uncultured tracheal aspirates were further subjected to species identification and “carbapenamase” genotyping by multiplex PCR.

## Materials and Methods

The study included 75 clinical isolates of *Acinetobacter baumannii* and 153 purulent tracheal aspirates were used for the development and validation of our assay. The clinical isolates and the tracheal aspirates, which formed part of routine work, were provided by the clinical microbiology laboratory at two tertiary care hospitals in Hyderabad. Species identification and antimicrobial susceptibility testing were done in the microbiology laboratories on VITEK-2 in accordance with CLSI guidelines and results were decoded later for comparative analysis with our CarbaNP and genotyping studies.

### Preparation of genomic DNA of *A. baumannii*

Pure single colony of each isolate was suspended in 100 μl of TEX buffer (10 mM Tris-HCl pH 8.5, 1 mM EDTA, 1% (w/v) Triton X100). The suspension was vortexed to achieve a uniform suspension and centrifuged. The pellet was subjected to another wash in TEX buffer. Finally the pellet was resuspended in 100 µl of TEX buffer and lysed by heating in a dry bath at 95°C for 15min [18]. The lysate was used as the DNA template for PCR amplifications. All isolates were screened by PCR using primers designed for the carbapenemase genes-*bla*_KPC_, *bla*_NDM-1_, *bla*_OXA-23-like_, *bla*_OXA-51-like_ and the species marker (Ab-ITS) of *A. baumannii* as described in “Table-1” [19-22]. Each selected target in this assay is the most common among carbapenemases described. The species marker (Ab-ITS), *bla*_OXA-23-like_, *bla*_OXA-51-like_ were identified by multiplex PCR and amplification conditions are as follows; initial denaturation at 95^0^C for 10min, followed by 35 cycles at 94^0^C for 30s, 50^0^C for 40s and 72^0^C for 50s and a final extension at 72^0^C for 5 min. Whereas *bla*_KPC_, *bla*_NDM_ PCR conditions consisted of an initial denaturation at 95^0^C for 10min, followed by 35 cycles of 94^0^C for 30s, 58^0^C for 40s, 72^0^C for 50s (*bla*_NDM_) and 94^0^C for 30s, 55^0^C for 30s (*bla*_KPC_) and 72^0^C for 50s, and a final extension at 72^0^C for 10min for both gene targets. The multiplex PCR products of 501bp (*bla*_OXA-51-like_), 353bp (*bla*_OXA-23-like_), 208bp (Ab-ITS) and amplicons of 660bp (*bla*_NDM-1_), 246bp (*bla*_KPC_) were visualized in a 2% agarose gel after electrophoresis (w/v) and staining with ethidium bromide.

**Table-1:**
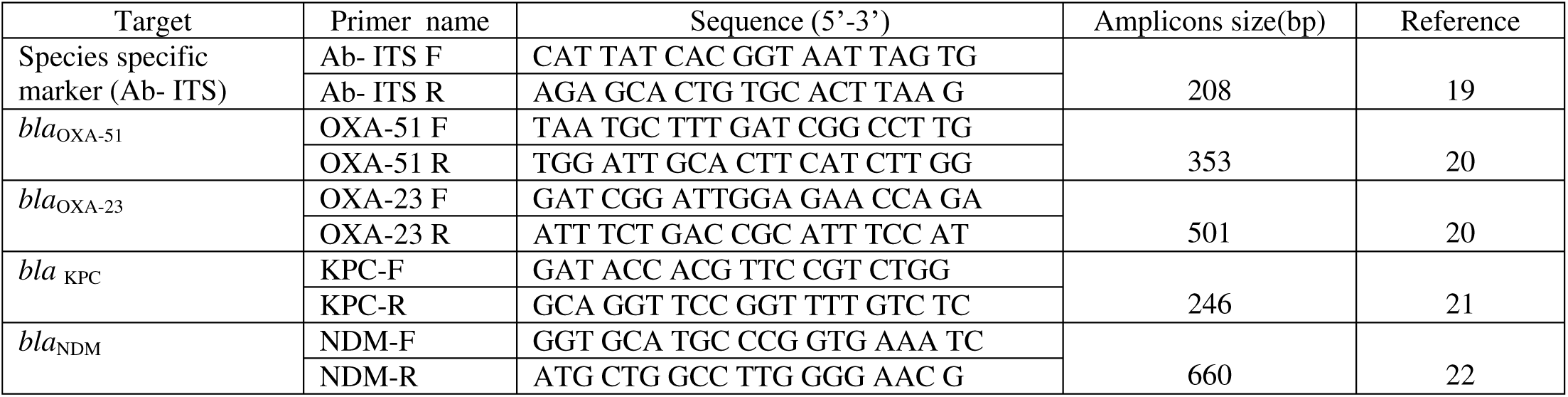
Primers used in this study:

### CarbaNP Assay

The CarbaNP assay (Nordmann *et al*, 2012) was performed as described below with some modifications to the original protocol. Colonies of *A. baumannii* grown on Muller Hinton Agar plates with imipenem disks for 24h were used. A single calibrated loop (5 μL) of bacterial mass was suspended in 100μL of TX buffer (10 mM Tris-HCl pH 8.5, 1% w/v Triton X-100) vortexed for one minute and further incubated at 37^0^C for 1hour. The lysed bacteria were centrifuged at 10,000 rpm at room temperature for 5 min and the cell free supernatant was used for carbapenamase assay. Tracheal aspirates were processed as follows (Figure-1): 150 μL of the tracheal aspirate was taken into 1.5mL eppendorf tube and centrifuged for 5 min at 10,000rpm. The supernant was discarded and the pellet was washed as before with TX buffer twice; finally 100 μL of TX lysis buffer was added to the pellet and subjected to vortex in pulses of 30 s for 2 min, incubated at 37^0^C for 1h for maximum lysis of bacteria and centrifuged for 5 min at 10,000rpm. The cell free supernatant was used for the CarbaNP assay. The assay was performed in sterile 96 well microtiter plate. 30 μL of cell free supernant was mixed with 100 μL assay solution containing 3mg/L of imipenem, 40 μL of 0.1 mM ZnSO4 and 30 μL of phenol red indicator (pH8.5) in respective wells. 5 μL of 3mM of Ethylene Diamine Tetra Acetic acid disodium salt (EDTA for Metallo β-lactameses) or 1 μL of 8 mg/ml of Phenyl Boronic Acid (PBA for Class A enzymes) or 1 μL of 100 mM of NaCl (for Class D enzymes) was added depending on which enzyme was being assayed. After addition of inhibitor, 40 μL of 0.1 mM ZnSO4 and 30 μL of phenol red (pH8.5) were added to the wells and incubated at 37^0^C up to 2 h. The carbapenemase activity was identified by a change in color of phenol red to yellow. The absorbance of the solution in the microtiter wells was measured at different time intervals (T_0_, T_15_, T_30_, T_60_ and T_120_ min) at 546 nm.

**Fig 1:**
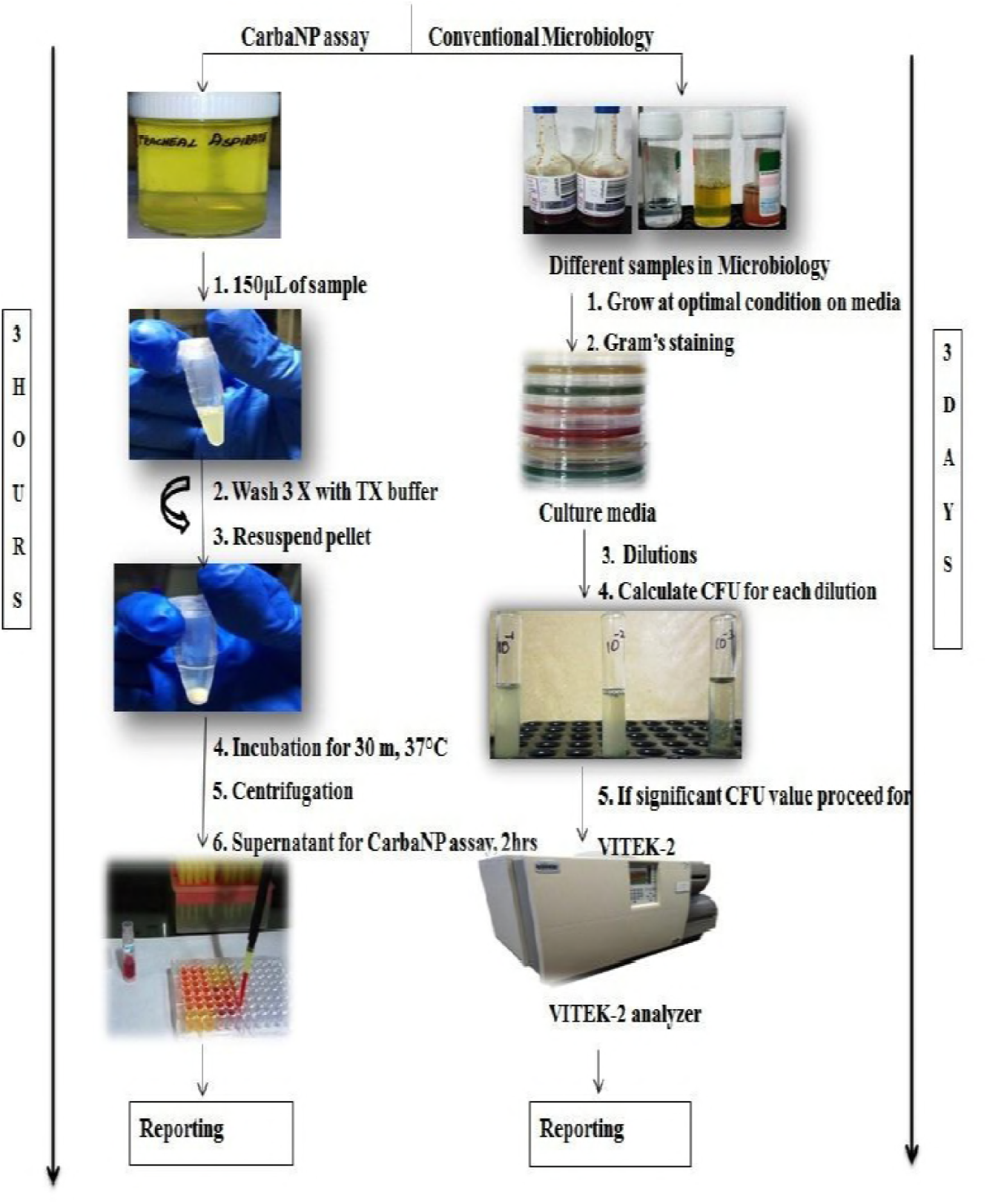
Flow diagram of processing of tracheal aspirates

## RESULTS

### Carbapenemase detection

To extend the chromogenic assay protocol to uncultured clinical samples: first we optimized the assay on a few culture isolates (n= 75) of *A. baumannii*. These isolates were screened for species specific marker Ab-ITS (16S-23S rRNA spacer region) and carbapenemase genes through PCR. The distribution of carbapenemase genes among these culture isolates of *A. baumannii* were as follows: most of them, 97.3% (73/75), carried the *bla*_OXA-51-like_ gene, and *bla*_OXA-23-like_ gene, 85.3% (64/75). Further, *bla*_NDM-1_ and *bla*_KPC_ were detected in 44% (33/75) 30.6% (23/75) of the isolates. A total of 153 purulent tracheal aspirates were directly processed for CarbaNP test and genotyping. The phenotypic (VITEK-2) data were blinded to us. Hence, we screened them first for the species genetic marker (Ab-ITS) and subsequently were subjected to carbapenemase genotyping. Out of the 153 purulent tracheal aspirate samples, 103 (65.1 %) were identified as *A. baumannii* by species specific marker. 52 (32.9%) of them carried *bla*_OXA-51like_ and 37 (23.4%) carried *bla*_OXA-23like_. *bla*_KPC_ and *bla*_NDM-1_ were present in 26 (17%) and 39(25.4%) tracheal aspirate samples respectively. Genotypic results are presented in Table-2.

**Table −2:** Results of genotyping and modified CarbaNP on both carbapenem resistant isolates and clinical samples.

| Sample | Species marker | Class A | Class B | Class D |  | Modified carbaNP assay |  |
| --- | --- | --- | --- | --- | --- | --- | --- |
|  | Ab-ITS | KPC | NDM-1 | OXA-51 | OXA-23 | CarbaNP | Inhibitor |
| Culture isolates (n=75) | 75 | 23 | 33 | 73 | 64 | 68 | 64 |
| purulent aspirates (n=153) | 103 | 26 | 39 | 52 | 37 | 85 | 48 |

### Validation of CarbaNP assay

To validate CarbaNP assay *Escherichia coli* (NCTC 10418), *Pseudomonas aeruginosa* (NCTC 10662), *Klebseilla pneumonia* (ATCC 700603), *Acinetobacter baumannii* (ATCC 19606) strains as positive controls and *Staphylococcus aureus* (ATCC 25923, ATCC 43300) strains were used as negative controls. The assay mixture contained buffered cell free extract, imipenem and ZnSO4 which when incubated at 37^0^C turned from red to orange or yellow depending on the enzyme activity or remained red for bacterial extracts which did not contain any carbapenemase. Our CarbaNP test results showed absolute correlation to our genotype results. Out of 75 culture isolates 73 contained *bla*_OXA-51,_ 24 contained *bla*_OXA-23_ plus *bla*_OXA-51_ and 3 were NDM producers (*bla*_OXA-51_ plus *bla*_NDM-1_). Multiple carbapenemase producers were 16 (*bla*_OXA-51_, *bla*_NDM-1_, *bla*_OXA-23_), 2 (*bla*_OXA-51_, *bla*_NDM_, *bla*_KPC_), 10 (*bla*_OXA-51_, *bla*_OXA-23_ and *bla*_KPC_) and 12 (*bla*_OXA-51_, *bla*_OXA-23_, *bla*_NDM_ and *bla*_KPC_) respectively. 6 isolates were completely negative for carbapenemases other than *bla*_OXA-51_. The rate of hydrolysis of imipenem was monitored at different time periods *i. e* 0 min, 15min, 30min, 60min and 120min with and without the appropriate carbapenamase inhibitor. The rate of hydrolysis was different for single carbapenemase, two carbapenemase and multiple carbapenemase producers. The results of these enzyme assays are represented in fig-2. EDTA was used as inhibitor for Class B (Metallo Beta Lactamase, MBL) type carbapenemases, NaCl for Class D (Oxacillinases, OXA) and Phenyl Boronic Acid (PBA) was used as specific inhibitor of Class A type (*K. pneumoniae* carbapenamase, KPC) enzymes. CarbaNP assay results “(see table 3)” were interpreted as follows: 1) if the color turned from red to orange or yellow in the presence of 3mM of EDTA (inhibits MBL) and 100mM of NaCl (inhibits OXA type) whereas wells containing PBA remained red in color, the isolate was KPC producer; 2) if the color in the wells turned from red to orange or yellow in the presence of PBA and 100mM of NaCl whereas wells containing EDTA remained red in color, the isolate was MBL (*bla*_NDM-1_) producer; 3) if the color in the wells turned from red to orange or yellow in the presence of PBA and EDTA whereas wells containing NaCl remained red in color, the isolate was oxacillanase (*bla*_OXA-23_ and *bla*_OXA-51_) producer; and 4) if the color of the wells remained red under all the above conditions, the isolate was considered a non-carbapenemase producer. Assay results were categorized based on carbapenemase genotype data into 4 groups: (i) all positive, (ii) OXA-51, OXA-23 positive and others negative, (iii) NDM negative and others positive and (iv) KPC negative and others positive. Colorimetric changes after 30 min incubation at 37^0^C are presented in fig-3. Absorbance at 546nm of all categories were noted and represented in graph. 30 min incubation was used for screening and for inhibitor assays. Results were depicted in fig-4.

**Table 3:** Modified CarbaNP results with inhibitors for different classes of carbapenemases

| S. No | Reaction mixture + Inhibitor | Color change | Enzyme category |
| --- | --- | --- | --- |
| 1 | 1. EDTA +NaCl | Red → Orange /Yellow | CLASS A |
|  | 2. PBA | Red (No change) |  |
| 2 | 1. PBA + NaCl | Red → Orange /Yellow | CLASS B |
|  | 2. EDTA | Red (No change) |  |
| 3 | 1. PBA + EDTA | Red → Orange /Yellow | CLASS D |
|  | 2. NaCl | Red (No change) |  |
| 4 | 1. EDTA +NaCl | Red (No change) | NO CARBAPENEMASE<br>ACTIVITY |
|  | 2. PBA + EDTA | Red (No change) |  |
|  | 3. PBA + NaCl | Red (No change) |  |

**Fig −2:**
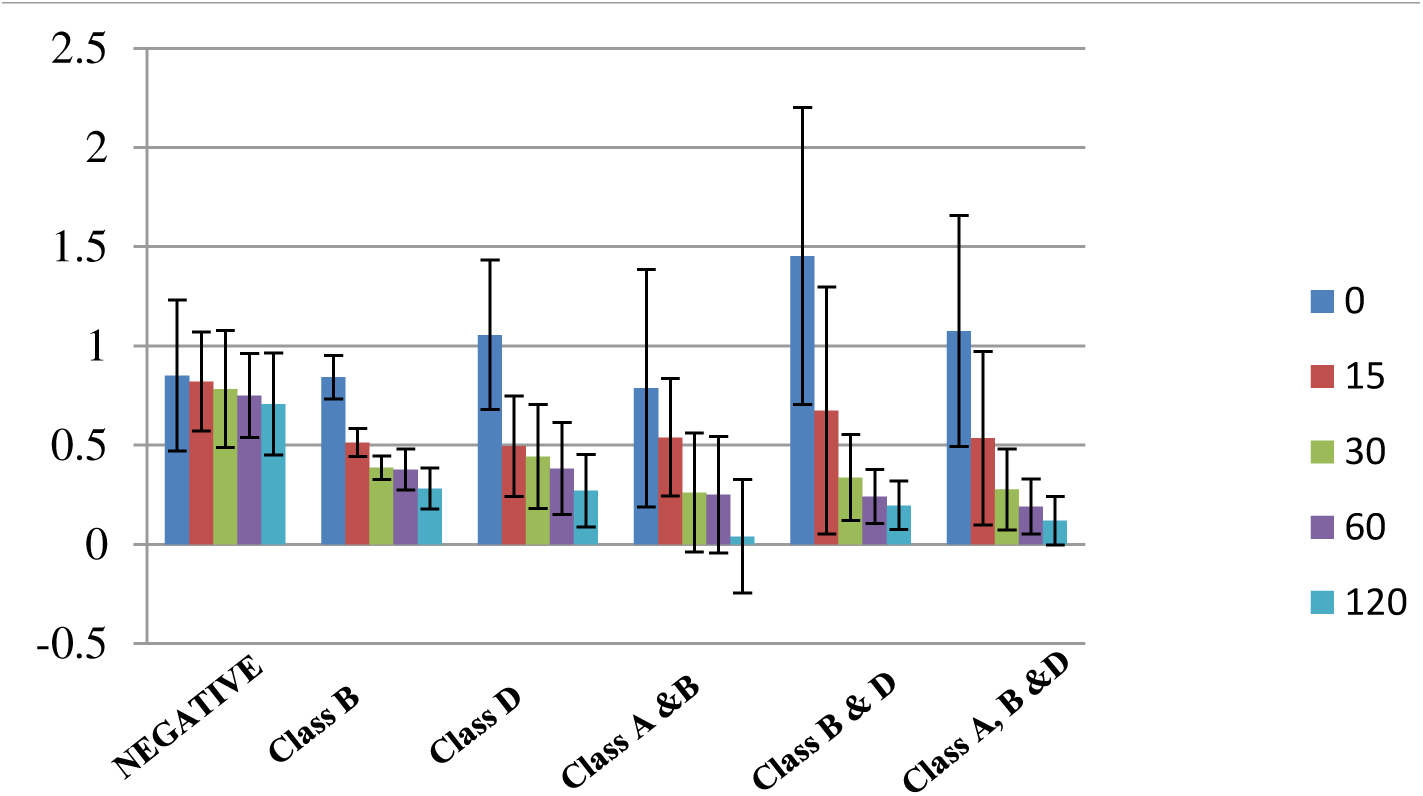
Rate of hydrolysis between single, double and multiple carbapenemase producers **Graphical representation of rate of hydrolysis at T_0_, T_15_, T_30_, T_60_ and T_120_**

1. Negative: non carbapenemase producers
2. Class B: *bla*_NDM_
3. Class D: *bla*_OXA-51_; *bla*_OXA-23_
4. Class A &B: *bla*_KPC_ *bla*_NDM_
5. Class B & D: *bla*_NDM_, *bla*_OXA-51_, *bla*_OXA-23_
6. Class A, B & D: *bla*_KPC_, *bla*_NDM_, *bla*_OXA-51_, *bla*_OXA-23_
7. Error bars represent standard deviation of calorimetric measurements at 546nm.

**Fig −3:**
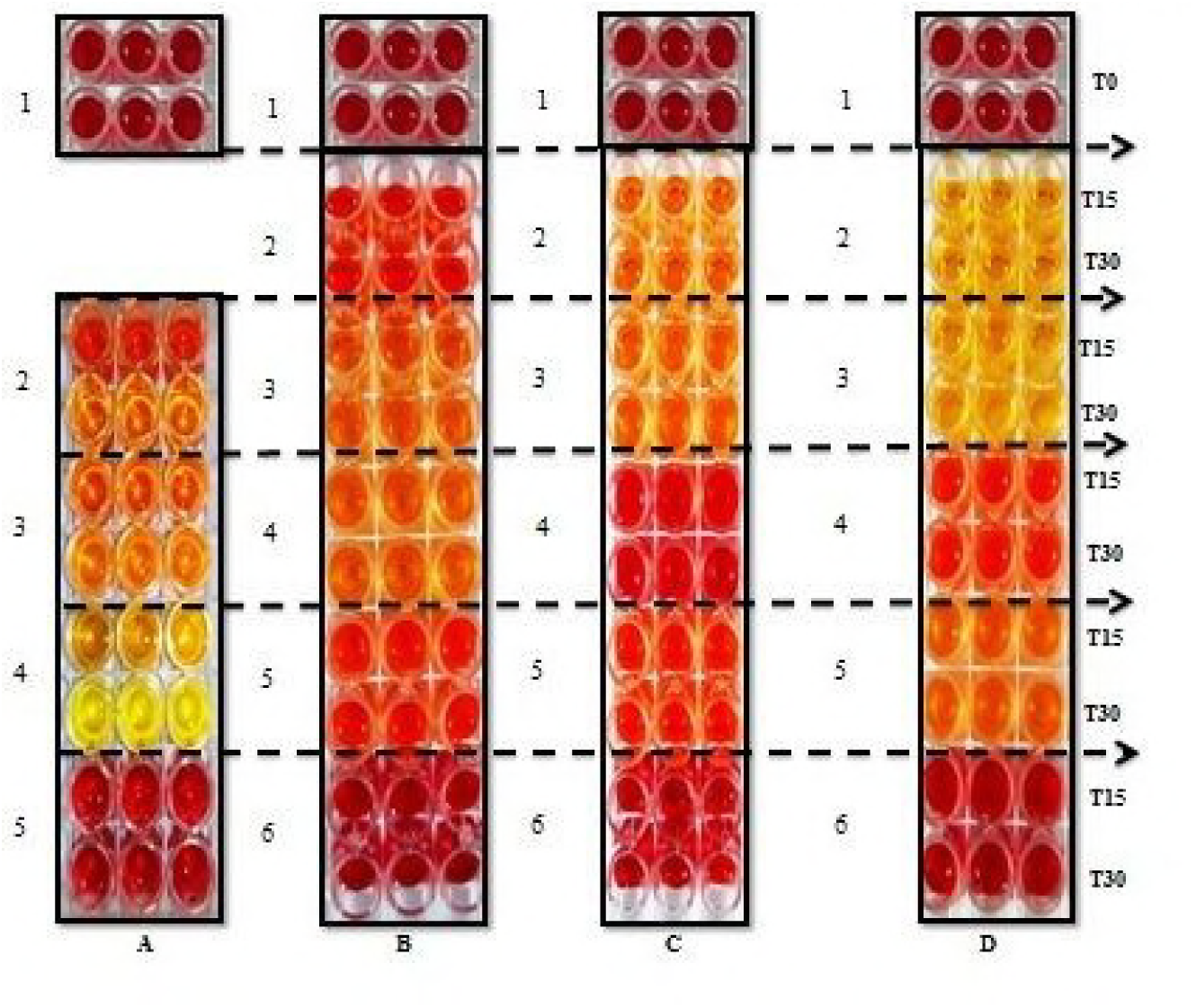
Colorimetric changes during modified carbaNPassay in clinical isolates of *A. baumanniii*. **A:ALL POSITIVE**; 1. Reaction mixture at T_0_, 2. Imipenem solution + ZnSO4+ PBA, 3. Imipenem solution + ZnSO4+ EDTA, 4. Imipenem solution + ZnSO4+ NaCl, 5. No imipenem **B: Class B Negative and others positive**; 1. Reaction mixture at T_0_, 2. Imipenem solution + ZnSO4+ PBA, 3. Imipenem solution + ZnSO4+ EDTA, 4. Imipenem solution + ZnSO4-EDTA 5. Imipenem solution + ZnSO4+ NaCl, 6. No imipenem. **C: Class D positive and others negative:** 1. Reaction mixture at T_0_, 2. Imipenem solution + ZnSO4+ PBA, 3. Imipenem solution + ZnSO4+ EDTA, 4. Imipenem solution + ZnSO4 + NaCl, 5. Imipenem solution + ZnSO4-NaCl, 6. No imipenem. **D: Class A negative and others positive:** 1. Reaction mixture at T_0_, 2. Imipenem solution + ZnSO4+ PBA, 3. Imipenem solution + ZnSO4 - PBA, 4. Imipenem solution + ZnSO4+ EDTA, 5. Imipenem solution + ZnSO4+ NaCl, 6. No imipenem.

**Fig-4:**
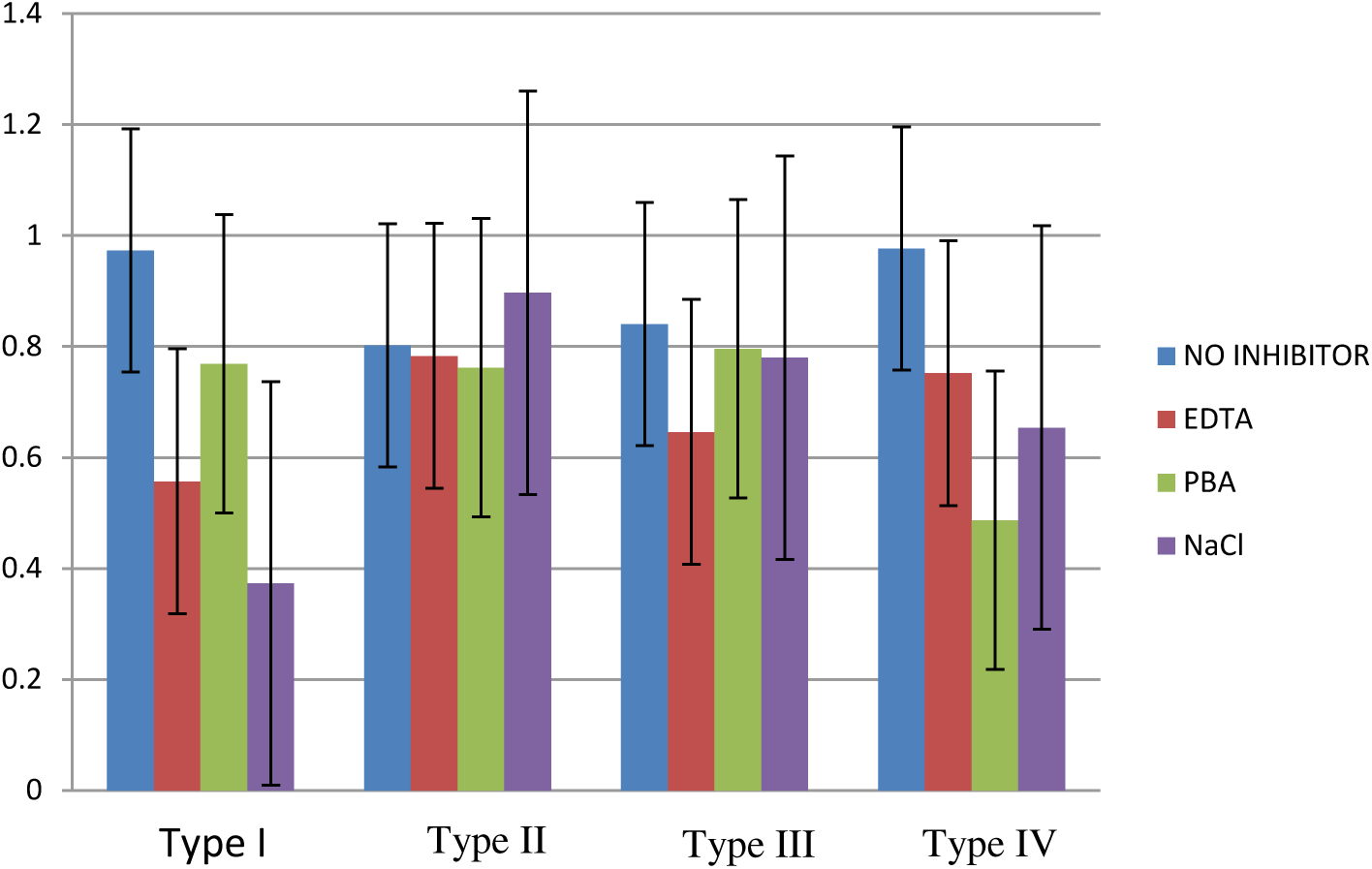
Rate of hydrolysis at T_30_ with inhibitors. **The carbapenemase activity and the effect of inhibitors.** The effect of inhibitors (Control, EDTA, PBA & NaCl) on carbapenem resistant isolates of *Acinetobacter baumannii*. The bacterial extract was incubated with different inhibitors at room temperature for 2hrs. The enzyme activity was assayed here for 30 mins for all 4 categories. They are Type I: Class A (*bla*_KPC_), B(*bla*_NDM_) and D(*bla*_OXA-51_, *bla*_OXA-23_) (+); Type II: Class A(*bla*_KPC_) (-) and B(*bla*_NDM_) and D(*bla*_OXA-51_, *bla*_OXA-23_) (+); Type III: Class A (*bla*_KPC_) (+) and B (*bla*_NDM_) and D(*bla*_OXA-51_, *bla*_OXA-23_) (-); and Type IV: Class B (*bla*_NDM_) (-) and A(*bla*_KPC_) and D(*bla*_OXA-51_, *bla*_OXA-23_) (+). Error bars represent standard deviation of triplicate measurements.

The relative orders of rate of imipenem hydrolysis (total carbapenem activity) by these groups of clinical isolates of *A.baumannii* were as follows:

Class A, B and D **(+) (all positive)** > **Class A (-) and** B, D **(+)** > **Class D (+) and** B, D **(-)** > **Class B (-) and** A, D **(+).**

This summary of correlation between CarbaNP test and genotyping results may need further validation in other laboratories and for other genotypes of carbapenemases.

A total of 153 tracheal aspirates were included in this study. Phenotypic data of these clinical samples was blinded to our laboratory and the results were decoded after genotyping and CarbaNP study. Subsequently modified chromogenic assay was performed on the cell free extracts from these tracheal aspirates. These samples were processed as described earlier. Results of CarbaNP assay were compared with both VITEK-2 results as well as our genotype PCR results.

## Discussion

The massive dissemination of carbapenemase producers among *Enterobacteriaceae* demands that methods to quickly determine the microbial species and also the drug sensitivity profile are available in the microbiology laboratory. High sensitivity and specificity together with a rapid workflow has become mandatory to devise therapeutic strategies to treat dangerous pathogens and to control their spread [11, 12]. The application of CarbaNP in this study shows that it is capable of detecting different carbapenemases in Gram negative pathogens allowing distinction between enzymatic and non-enzymatic mechanisms of carbapenem resistance. The adaptation of CarbaNP method showed multiple advantages over the other phenotypic screening methods. CarbaNP has high sensitivity, specificity and is easily available for prompt identification of carbapenemases [11, 23]. Original CarbaNP test was validated on clinical isolates of various *Enterobacterial species* [2] and subsequently on clinical isolates of *A. baumannii* [17]. Recently the assay was adopted to detect “carbapenamase” activity in spiked blood cultures [24]. This method eliminates false positives compared to boronic acid synergy test, the modified Hodge test (MHT), and drastically reduces time compared to disc diffusion/E-Test methods [25, 26]. The carbapenem inactivation method (CIM), and the modified carbapenem inactivation method (mCIM) reported recently require at least 24-72h for results even though it costs less than <1$ [5, 27]. Rapidec CarbaNP test kit (BioMerieux, France), Neo-Rapid Carb screen kit, Rapid Carb Blue screen (ROSCO Diagnostics Denmark) are commercial kits available for screening clinical isolates for carbapenamase activity [14, 28, 29]. MALDI-TOF MS is effective method compared to Rapidec CarbaNP test kit to discriminate between carbapenemase and non-carbapenemase producers [30, 31]. As an alternative to molecular assay, antibody based methods such as lateral flow immunoassay (LFIA) has been developed for detection carbapenemase production. Lateral flow assay yield results from cultured strains within 15min with 100% sensitivity and specificity [32, 33]. The manual versions of rapid calorimetric assays includes manual Carba NP CLSI method, manual Blue Carba, and modified Carba NP which requires reagent preparation compared to ready to use kit methods [16,29, 34]. These CarbaNP tests available till date help to reduce the time taken to identify carbapenamase activity in clinical isolates. However, the full potential of the test could be realised only when the test is applicable directly to uncultured clinical samples so that therapeutic decisions can be made quickly. We attempted to fulfill this gap. We have simplified the cell free extract preparation and validated it directly on purulent tracheal aspirates to detect, quantitate and also differentially demonstrate various carbapenamase activities using specific inhibitors. Remarkable correlation was observed in our study between the genotypic test results (PCR) and CarbaNP assay results. CarbaNP method greatly reduces cost and labor with excellent accuracy in results even with uncultured tracheal aspirates. The convenience of CarbaNP test is notable as it doesn’t require any specialized equipment and needs a short hands-on time to perform and requires only single colony from the agar plate or 150 µL of the tracheal aspirate. It is easily scalable to a high throughput format for large number of Gram negative isolates and adaptable in many laboratories. This adapted assay can be used for rapid differential diagnosis of carbapenemases directly in clinical samples and clinical isolates. Blindfolding the tracheal aspirate (n=153) samples for the genotyping/CarbaNP screening by the microbiology laboratory shows that our CarbaNP test format is applicable and adaptable as a routine in a clinical microbiology laboratory. Remarkable correlation between genotyping and CarbaNP test results shows that this colorimetric assay is reliable and accurate. We have optimized the average incubation time as 30 min. Inclusion of inhibitors seem to be an attractive option to narrow down the drug sensitivity profile compared to expensive, time consuming disc diffusion or E-test. *bla*_KPC_, *bla*_NDM_ were used as typical member of the Class A and B enzymes in PCR genotyping. Considering the prevalence in *A. baumannii,* both *bla*_OXA-23_ and *bla*_OXA-51_ were screened for Class D (CHD) in culture isolates. Whereas in tracheal aspirates Class D enzymes (*bla*_OXA-23_ and *bla*_OXA-51_), Class B (*bla*_NDM_) and Class A (*bla*_KPC_) were screened to correlate with phenotyping results along with species identification. The data from these genotyping were used to evaluate the merit of the CarbaNP test for differentiating various classes of carbapenemases. We introduced certain modifications to Nordmann’s protocol to make it more user friendly, to reduce the quantity of cells required and to apply it directly to uncultured tracheal aspirates. These modifications include (i) extraction buffer i.e TX buffer (10 mM Tris-HCl pH 8.5, Triton X 100, 1% w/v) used for bacterial cell lysis (ii) Extraction was performed by vigorous vortexing and incubation at 37^0^C for one hour for cell lysis (iii) Rate of imipenem hydrolysis was monitored at 546 nm for different time periods: 0m, 15m, 30m, 60m and 120m. This assay clearly differentiated various classes of carbapenemases in culture isolates and uncultured clinical samples. We are validating our protocol on other types of clinical samples currently and also for other gram negative pathogens. All the carbapenem resistant isolates showed some (intrinsic) carbapenemase activity due to the presence of *bla*_OXA-51_ and other isolates showed incrementally higher hydrolysis depending on whether they contained one, two or more carbapenemases. Whereas reaction is completely inhibited in the presence of inhibitors resulting in no change in the absorbance at 546 nm at different time periods. Graphical representation of rate of hydrolysis with or without inhibitors clearly indicates that identification and differentiation of carbapenemase activity would be a valuable tool in characterization of isolates and monitoring their spread. The sensitivity and specificity of this assay were found to be excellent when compared with VITEK-2 (sensitivity and specificity was 100%). This CarbaNP test offers an economical and faster way to profile the carbapenemases in isolates of *A. baumannii* and also in uncultured tracheal aspirates faster than even molecular techniques. The format can be easily adopted for a high throughput screening of carbapenemases in clinical samples or isolates. We hope it will become a valuable and useful tool in clinical microbiology screening and to study the epidemiology of carbapenemase producers [12].

## Conclusion

We propose the following algorithm to screen uncultured clinical samples directly for carbapenemase activity:

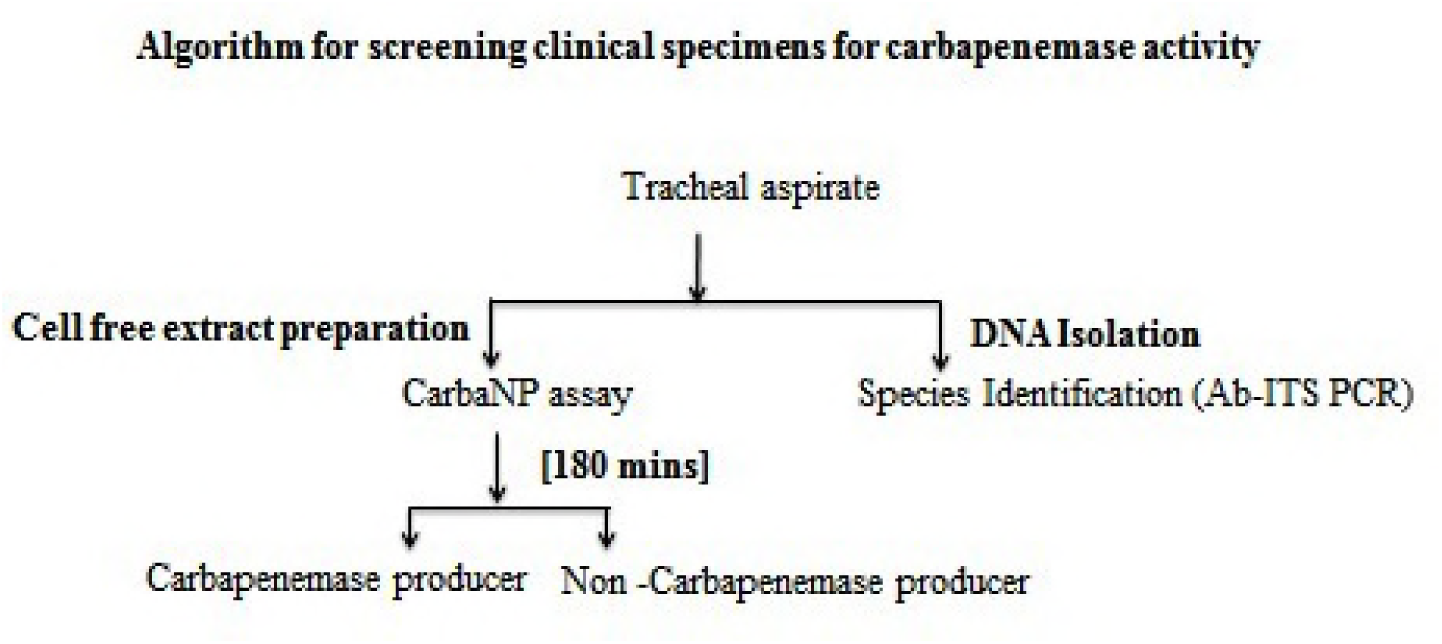

This algorithm will intensify the quick identification of carbapenemases for proper infection control measures. The conventional microbiology results are made available in 3-4 days for species identification and drug sensitivity profile. Our protocol which includes a simple one step PCR for species identification and CarbaNP test determination of carbapenamase activity promises to make the same information available to the clinician the same day, both together precisely in less than 4 h. This is potentially a remarkable improvement and it is hoped that will revolutionalise the way multidrug resistance will be identified and used for the management of patients and to control its spread in the society.

## Acknowledgement

This work was supported by Global Medical Education Research and Foundation (GMERF), Hyderabad.

## References

1. Nordmann P, Poirel L. 2013. Strategies for identification of carbapenemase-producing *Enterobacteriaceae*. J. Antimicrob. Chemother. 68:487–489.

2. Nordmann P, Poirel L, Dortet L. 2012. Rapid detection of carbapenemase-producing *Enterobacteriaceae*. Emerg. Infect. Dis. 18:1503–1507.

3. Osterblad M, Hakanen AJ, Jalava J. 2014. Evaluation of the Carba NP test for carbapenemase detection. Antimicrob. Agents Chemother. 58:7553–7556.

4. Nordmann P, Naas T, Poirel L. 2011. Global spread of carbapenemase producing *Enterobacteriaceae*. Emerg. Infect. Dis. 17:1791–1798.

5. Van der Zwaluw K, de Haan A, Pluister GN, Bootsma HJ, de Neeling AJ, Schouls LM. 2015. The carbapenem inactivation method (CIM), a simple and low-cost alternative for the Carba NP test to assess phenotypic carbapenemase activity in gram-negative rods. PLoS One 10:e0123690.

6. Livermore DM. 2001. Of *Pseudomonas*, porins, pumps and carbapenems. J. Antimicrob. Chemother. 47: 247–250.

7. Jocoby GA, Mills DM, Chow N. 2004. Role of beta-lactamases and porins in resistance to ertapenem and other beta-lactams in *Klebsiella pneumoniae*. Antimicrob. Agents Chemother. 48: 3203–3206.

8. Carattoli A. 2013. Plasmids and the spread of resistance. Int J Med Microbiol. 303: 298–304.

9. Nordmann P, et al. 2012. Identification and screening of carbapenemaseproducing *Enterobacteriaceae*. Clin. Microbiol. Infect. 18:432–438.

10. Lee LY, Korman TM, Graham M. 2014. Rapid time to results and high sensitivity of the CarbaNP test on early cultures. J. Clin. Microbiol. 52:4023–4026.

11. Dortet L, Poirel L, Nordmann P. 2012. Rapid detection of carbapenemase-producing *Pseudomonas spp*. J. Clin. Microbiol. 50:3773–3776.

12. Dortet L, Poirel L, Nordmann P. 2012. Rapid identification of carbapenemase types in *Enteriobacteriaceae* and *Pseudomonas spp.* by using a biochemical test. Antimicrob. Agents Chemother. 56: 6437–40.

13. Tijet N, Boyd D, Patel SN, Mulvey MR, Melano RG. 2013. Evaluation of the Carba NP test for rapid detection of carbapenemase-producing *Enterobacteriaceae* and *Pseudomonas aeruginosa.* Antimicrob. Agents Chemother. 57:4578–4580.

14. Hombach M, Von Gunten B, Castelberg C, Bloemberg GV. 2015. Evaluation of the Rapidec Carba NP test for detection of carbapenemases in *Enteriobacteriaceae*. J. Clin. Microbiol. 53: 3828–33.

15. Queenan AM, Bush K. 2007. Carbapenemases: the versatile beta-lactamases. Clin. Microbiol. Rev 20: 440–458.

16. J. Pires, A. Novais and L. Peixe.2013. Blue-Carba, an easy biochemical test for detection of diverse carbapenemase producers directly from bacterial cultures. J.Clin. Microbiol. 51: 4281–4283.

17. Dortet L, Poirel L, Errera C, Nordmann P. 2014. CarbAcineto NP test for rapid detection of carbapenemase producing *Acinetobacter spp*. J. Clin. Microbiol. 52: 2359–2364.

18. Sritharan V and Barker RH. 1991. A simple method for diagnosing *M. tuberculosis* infection in clinical samples using PCR. Mol. cell probes. 5: 385–395.

19. Chen TL, Siu LK, Wu RC, Shaio MF, Huang LY, Fung CP. 2007. Comparison of one-tube multiplex PCR, automated ribotyping and intergenic spacer (ITS) sequencing for rapid identification of *Acinetobacter baumannii*. Clin. Microbiol. Infect. 13 : 801–806.

20. Amudhan SM, Sekar U, Arunagiri, Sekar B. 2011. OXA beta-lactamase –mediated carbapenem resistance in *Acinetobacter baumannii*. Indian J. Med. Microbiol. 29 : 269–274.

21. Hindiyeh M, Smollen G, Grossman Z, Ram D, Davidson Y, Mileguir F. 2008. Rapid detection of _*bla*KPC_ carbapenemase genes by Real-Time PCR. J. Clin. Microbiol. 46:2879–2883.

22. Bonnin RA, Naas T, Poirel L, Nordmann P. 2012. Phenotypic, Biochemical and molecular techniques for detection of metalo-beta-lactamase NDM in *Acinetobacter baumannii*. J. Clin. Microbiol. 50: 1419–1421.

23. Vasoo S, Cunningham SA, Kohner PC, Simner PJ, Mandrekar JN, Lolans K, Hayden MK, Patel R. 2013. Comparsion of a novel, rapid chromogenic biochemical assay, the CarbaNP test, with the modified Hodge test for detection of carbapenemase producing Gram negative bacilli. J. Clin. Microbiol. 51:3097–3101.

24. Dortet L, Brechard L, Poirel L, Nordmann P. 2014. Rapid detection of carbapenemase producing *Enterobacteriaceae* from blood cultures. Clin. Microbiol. Infect. 20: 340–344.

25. Doi Y, Potoski BA, Adams-Haduch JM, Sidjabat HE, Pasculle AW, Paterson DL. 2008. Simple disk-based method for detection of *Klebsiella pneumoniae* carbapenemase-type beta-lactamase by use of a boronic acid compound. J.Clin.Microbiol. 46:4083–4086.

26. Carvalhaes CG, Picao RC, Nicoletti AG, Xavier DE, Gales AC. 2010. Cloverleaf test (modified Hodge test) for detecting carbapenemase production in *Klebsiella pneumoniae*: be aware of false positive results. J. Antimicrob. Chemother. 65:249–251.

27. Pierce VM, Simner PJ, Lonsway DR, Roe-Carpenter DE, Jhonson JK, Brasso WB. 2017.Modified Carbapenem Inactivation Method for phenotypic detection of carbapenem ase production among *Enterobacteriaceae*. J.Clin.Microbiol.55:2321–2333.

28. Garg A, Garg J, Upadhyay GC, Agarwal A, Bhattacharjee A. 2015. Evaluation of the Rapidec CarbaNP test kit for detection of carbapenemase producing gram negative bacteria. Antimicrob. Agents Chemother. 59: 7870–7872.

29. Tamma PD, Opene BN, Gluck A, Chambers KK, Carroll KC, Simner PJ. 2017. Comparison of 11 phenotypic assays for accurate detection of carbapenemase-producing *Enterobacteriaceae*. J.Clin.Microbiol.55:1046–1055.

30. Papagiannitsis CC, Studentova V, Izdebski R, Oikonomou O, Pfeifer Y, Petinaki E, Hrabak. 2015. Matrix-assisted laser desorption ionization-time of flight mass spectrometry meropenem hydrolysis assay with NH4HCO3, a reliable tool for direct detection of carbapenemase activity. J.Clin.Microbiol 53:1731–1735.

31. Choquet M, Guiheneuf R, Castelain S, Cattoir V, Auzou M, Pluquet E. 2018. Comparison of MALDI-TOF MS with the Rapidec Carba NP test for the detection of carbapenemase-producing *Enterobacteriaceae*. Eur. J. Clin. Microbiol. Infect. Dis. 37: 149–155.

32. Boutal H, Naas T, Devilliers K, Oueslati S, Dortet L, Bernabeu S, Simon S, Volland H. 2017. Development and validation of lateral flow Immunoassay for rapid detection of NDM-producing *Enterobacteriaceae*. J. Clin.Microbiol. 55: 2018–2029.

33. Boutal H, Voqel A, Bernabeu S, Devilliers K, Creton E, Cotellon E, et al. 2018. A multiplex lateral flow immunoassay for the rapid identification of NDM-, KPC-, IMP- and VIM-type and OXA-48-like carbapenemase-producing *Enterobacteriaceae*.J. Antimicrob. Chemother. 73: 909–915.

34. Bakour S, Garcia V, Loucif L, Brunel JM, Gharout-Sait A, Touati A, Rolain JM. 2015. Rapid identification of carbapenemase-producing *Enterobacteriaceae, Pseudomonas aeruginosa* and *Acinetobacter baumannii* using a modified Carba NP test. New. Microbes. New. Infect 7:89–93.

